# Insulin exposed endometrial epithelial cells cultured in a microfluidic device alters transcripts involved in translation that may contribute to reduced implantation capacity of the endometrium

**DOI:** 10.1101/2021.12.15.472777

**Authors:** Soo Young Baik, Haidee Tinning, Dapeng Wang, Niamh Forde

**Affiliations:** Discovery and Translational Sciences Department, Leeds Institute of Cardiovascular and Metabolic Medicine, Faculty of Medicine and Health, University of Leeds, LS2 9JT, United Kingdom; LeedsOmics, University of Leeds, Leeds, LS2 9JT, United Kingdom; Wellcome Centre for Human Genetics, University of Oxford, Oxford, OX3 7BN, United Kingdom

## Abstract

Obesity is a rapidly growing public health issue among women of reproductive age. It is also associated with decreased reproductive function including implantation failure. Implantation failure can result from a myriad of factors including impaired gametes and endometrial dysfunction. The mechanisms of how obesity-related hyperinsulinaemia disrupts endometrial function and implantation are poorly understood. Our study aims to investigate potential mechanisms by which insulin alters endometrial transcript expression, which may affect endometrial receptivity. Ishikawa cells mimicking human endometrial epithelium were seeded into a microfluidics organ-on-chip device to produce an *in vitro* endometrium. Syringe pump was attached to the microfluidics device to deliver three varying treatments into Ishikawa cells: 1) media control 2) vehicle control (PBS acidified to pH3 with acetic acid) 3) Insulin (2mg/mL) at a constant flow rate of 1uL/min for 24 hours to mimic secretion *in vivo*. Three biological replicates were obtained. Insulin-induced transcriptomic response of the *in vitro* endometrium was quantified via RNA sequencing, and subsequently analysed using DAVID and Webgestalt to identify Gene Ontology (GO) terms and signalling pathways. A Total of 29 transcripts showed differential expression levels across two comparison groups (control v vehicle control; vehicle control v insulin). There were nine transcripts significantly differentially expressed in vehicle control v insulin group (p<0.05). Functional annotation analysis of transcripts altered by insulin (n=9) identified three significantly enriched GO terms: SRP-dependent cotranslational protein targeting to membrane, poly(A) binding, and RNA binding (p<0.05). Over-representation analysis found three significantly enriched signalling pathways relating to insulin-induced transcriptomic response: protein export, glutathione metabolism, and ribosome pathways (p<0.05). Insulin-induced dysregulation of biological functions and pathways highlight potential mechanisms by which high insulin concentrations within maternal circulation may perturb endometrial receptivity.

## INTRODUCTION

Obesity is a complex disease with multifactorial aetiologies and in the UK, almost half of women within the childbearing age range are overweight or obese. Due to complex biochemical changes in the endocrinological and metabolic profile of those suffering from obesity, there are resulting complex health issues including insulin resistance and compensatory hyperinsulinaemia [1]. This has been linked to poor reproductive outcomes in women of childbearing age such as miscarriage, anovulation, irregular menstrual cycles, decreased conception rates in ART [2, 3].

The impact of obesity on ovarian function dysregulation is well recognised although complex. The higher concentrations of insulin in circulation associated with obesity can inhibit sex hormone binding globulin (SHBG) production by the liver [4] along with aromatisation of androgens to oestrogens associated with increased adipose tissue. As SHBG binds to circulating sex hormones; the combined effects of high peripheral aromatisation of androgens and decreased SHBG leads to an overall increase in bioavailable oestradiol and testosterone, therefore contributing to hyperandrogenism in the theca cells of the ovarian follicles [1] [2]. Hyperinsulinaemia and subsequent hyperandrogenism coupled with altered hormonal milieu can lead to premature follicular atresia and anovulation [2]. Furthermore, increased insulin in circulation associated with obesity can lead to inhibition of hepatic and ovarian IGF binding protein 1 (IGFBP1) expression an important regulator of ovarian and endometrial function. Overall, the combined effects of systemic insulin resistance and hyperinsulinaemia in obesity contribute to biochemical and sex hormone dysregulation within the female reproductive system. Such symptoms common features of polycystic ovarian syndrome (PCOS), and obesity related hyperinsulinaemia seems to exacerbate PCOS symptoms [5].

The effects that this dysregulation in metabolic and biochemical profiles associated with obesity have on endometrial function and uterine receptivity remains unclear. Conflicting results and discrepancies in methodologies between the studies may explain in part why this is the case. A series of studies from ovum donation cycles identified significantly higher spontaneous abortion rates in obese women (38.1%) compared to women with normal BMI (13.3%) [6], with a follow up study of 2656 first ovum donation cycles concluded that high BMI altered endometrial environment and function, although these effects are quite small [7]. A further investigation into 9,587 first cycle ovum donations showed that implantation, clinical pregnancy, and live-birth rates were significantly reduced in obese women, although miscarriage rates showed no difference with BMI [8] similar to what was observed by Cano et al, [9]. In contrast, no significant differences in receptivity impairment between obese and non-obese women were observed [10] or implantation rate or miscarriage rate in high BMI groups compared to lower BMI [11] although in this study miscarriage rates that were disproportionately high.

The studies critiqued above illustrate the controversial nature of the effect of obesity on endometrial function/implantation using the ovum donation model to isolate the impact of endometrial factors from the embryonic influence in implantation capability. This model limits study participants only to those participating in oocyte donation to remove the influence of poor embryo quality on implantation failure. Given there are likely fundamental differences in those that require ovum donation in obese and non-obese scenarios these are likely to. Compound results [12, 13]. Overall, these clinical studies highlight the gap in knowledge in regard to the mechanism by which the metabolic and biochemical changes in circulation contributes to dysfunction in endometrial function and receptivity.

Glucose metabolism is important during the peri-implantation window with expression of the facilitative glucose transporters (GLUT) in endometrial epithelia modulated by oestradiol and progesterone [14]. Furthermore, upregulation of glucose and its transporter (*GLUT1*) are critical for endometrial stromal cell decidualisation [15]. Obesity-related insulin resistance and hyperinsulinaemia leads to disturbances in glucose metabolism, which may have downstream effects on endometrial function and receptivity [16]. Dissecting out the complex interactions between insulin-specific alterations on endometrial function without the confounding influence of additional metabolic stressors in circulation it difficult. Moreover, static culture systems *in vitro* do not recapitulate exposure *in vivo*. Evidence in other species looking at the proteomic analysis on endometria of sows with normal reproductive performance and low reproductive performance to pinpoint the differentially expressed genes and pathways (ie. Insulin-signalling pathway and lipid metabolism) associated with uterine impairment. Chen et al. subsequently used insulin-resistant animal models driven by high-fat diet and Ishikawa cells (human endometrial adenocarcinoma cells) to create *in vitro* implantation models and reported that insulin resistance reduces receptivity through mitochondrial dysfunction and consequent oxidative stress [17]. Additionally, the use of a microfluidics endometrium-on-a-chip in bovine demonstrated cellular transcriptional and secretome differences if the endometrium following exposure to physiological extremes of metabolic factors glucose and insulin in microfluidics [18].

However, both models used animal endometrial tissue, and these findings can only be applied to humans to some extent. Various research investigating whether obesity impairs endometrial receptivity reveals conflicting findings; they also lack validity due to methodological limitations and complications associated with using the ovum-donation model. There is an overall scarcity in literature highlighting hyperinsulinaemia in obesity as an aetiology of poor uterine function and implantation. We therefore propose to investigate the impact of obesity-related metabolic stressor of insulin on endometrial function. To achieve this, we used a microfluidics approach to better mimic *in vitro*, exposure of the endometrial epithelium to Insulin that would occur *in vivo*. This allowed us to identify the transcriptional response of the endometrial epithelium revealing potential mechanism by which hyperinsulinaemia in obesity may contribute to endometrial dysfunction.

## MATERIALS AND METHODS

Unless otherwise stated all consumables were sourced from Sigma Aldrich (UK).

### Culture of Ishikawa cells and device preparation

Passage 20-24 Ishikawa cells (Immortalised human endometrial cells; ECACC #99040201) cultured in DMEM/F-12 [Dulbecco’s Modified Eagle Medium/Nutrient Mixture F-12], containing 10% FBS [foetal bovine serum], 1% GSP [glutamine, streptomycin, penicillin]) were trypsinised from a T75 flask and counted using a hematocytometer. Media (described above) was added to cells until they reached a concentration of 1,000,000/mL. The microfluidics device (Ibidi u-slide V0.4) were placed in the incubator at 37°C. One hundred microlitres of the of 1,000,000/mL Ishikawa cell suspension was subsequently seeded into each channel of V0.4 slides, and left to attach for 12 hr. Media (DMEM/F12, 10% FBS, 1% GSP) was added to each channel until almost full to prevent channel from drying. Ishikawa cells were left for 24 hr in 37°C incubator with 5% CO_2_ before undergoing treatment in the microfluidics device.

### Treatment of Ishikawa cells with Insulin in a microfluidic device

Each channel of the microfluidics device was flushed five times with 37°C PBS and the inlet syringes were connected with sterile tubing. Elbow connectors were then used to attach the sterile tubing the microfluidics device inlet. Sterile tubing was filled with medium from the syringes and attached to the device. A droplet to droplet method was used to prevent introduction of bubbles into the device. Syringes were pushed to fill both the chamber and outlet. The outlet tubing was filled with sterile PBS and connected to the outlets on microfluidics channels using the droplet-droplet method. The other end of each outlet tube was connected to a 7mL bijou container to collect and save the conditioned medium. After setting up the microfluidics device, medium was pushed through the entire device system to check for bubbles, and to get a droplet at the end of each outlet tube inside the bijou container. The microfluidics device set-up can be pictured in Figure 1.

**Figure 1.**
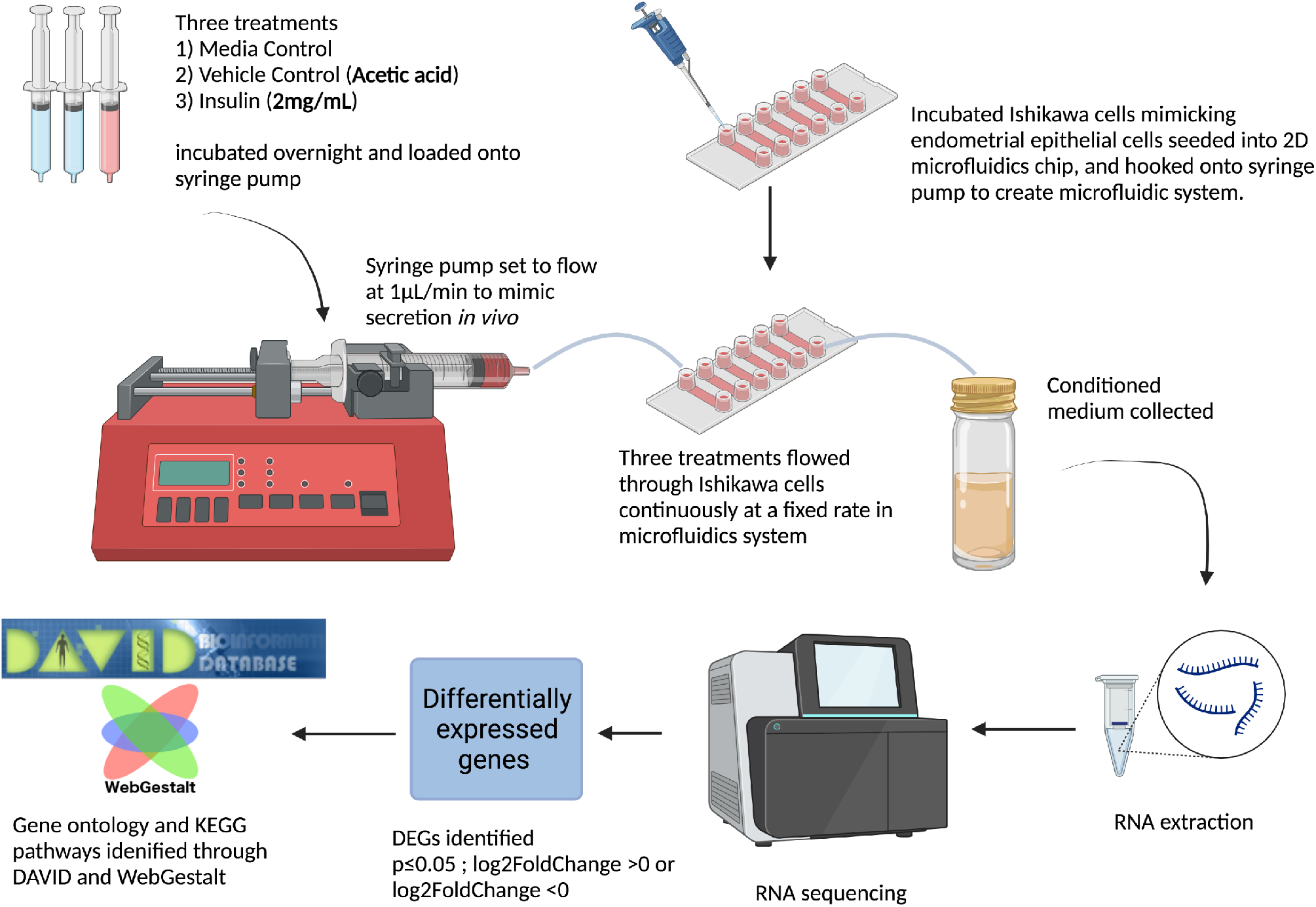
Schematic diagram of experimental setup and work flow of experiment. Diagrammatic representation of experimental approach using a microfluidics approach to mimic exposure to endometrial epithelial cells (n=3 biological replicates) to media control, vehicle control, or Insulin (2ml/mL) for 24 hr. Cells were analysed for a transcriptional response via RNA sequencing. Images produced with Biorender.

Cells within the devices were exposed to one of following three treatments for 24 hr (n=3 biological replicates): 1) media control, 2) vehicle control (PBS [phosphate-buffered saline] acidified to pH3 with acetic acid [A6283]), and 3) Insulin (2 mg/ml in PBS acidified to pH3 with acetic acid). Each treatment was loaded into a 5mL syringe and put in 37°C incubator overnight and loaded onto the syringe pump on the day of microfluidics run. The syringe pump and microfluidics system were placed into a 37°C incubator and the syringe pump was set to flow at the rate of 1μL/min in order to mimic secretion *in vivo* (as previously described; [18]).

### Cell Recovery and RNA extraction and RNA sequencing

After disconnecting the microfluidics device from the system, every channel from the device was flushed with PBS. Ishikawa cells were lifted with trypsin, neutralised with media, and centrifuged at 500g for 5 min to pellet the cells. The resulting cell pellet was then snap frozen in liquid nitrogen until RNA extraction. RNA extraction was carried by using the Mini RNeasy Kit (Qiagen), according to instructions provided by manufacturer. Extracted RNA was sent to Novogene (Cambridge, UK) for subsequent library preparation. Ribosomal RNA was firstly removed and a directional sequencing library was constructed using NEBNext® UltraTM Directional RNA Library Prep Kit for Illumina® (NEB, USA) following manufacturer’s protocol. Indices were included to multiplex multiple samples. Briefly, the first strand cDNA was synthesized using random hexamer primers followed by the second strand cDNA synthesis. The strand-specific library was ready after end repair, A-tailing, adapter ligation, size selection, and USER enzyme digestion. After amplification and purification, insert size of the library was validated on an Agilent 2100 and quantified using quantitative PCR (Q-PCR). Libraries were then sequenced on Illumina NovaSeq 6000 S4 flowcell with PE150 according to results from library quality control and expected data volume.

### Bioinformatic analysis

The raw and processed FASTQ files were scanned for the fundamental quality control metrics using FastQC (https://www.bioinformatics.babraham.ac.uk/projects/fastqc/). The adapter sequences in the reads were trimmed off using Cutadapt [19] and poor quality bases or reads were trimmed or discarded using PRINSEQ [20]. The human reference genome file (GRCh38) and corresponding GTF file were downloaded from GENCODE Human Release 31 [21]. The processed paired-end reads were aligned onto the human reference genome using STAR [22]. The resultant alignment files were converted, sorted and indexed with SAMtools [23] and only uniquely mapped reads were selected for the downstream analysis. The reads were counted for each gene using featureCounts() function of Rsubread package and a matrix of raw read counts for all genes was produced [24]. The differential expression testing was performed using DESeq2 with apeglm method for the log2 fold change shrinkage > 0.1 (or < - 0.1) and p value of < 0.05 [25].

Differentially expressed protein-coding genes found in three treatment groups were subjected to Gene Ontology (GO) functional annotations and over-representation enrichment analysis using DAVID (The Database for Annotation, Visualization and Integrated Discovery; Bioinformatics Resourced 6.8 [26]) and WebGestalt (WEB-based Gene SeT AnaLysis Toolkit [27]), respectively. Gene identifier for DAVID was chosen as “OFFICIAL_GENE_SYMBOL,” with list type selected as “Gene List.” When using WebGestalt, the functional database was chosen as “KEGG Pathway,” and gene ID type was “Gene symbol.” Gene reference set when using WebGestalt was “genome protein-coding.” P-value cutoff was p<0.05. P-values for both DAVID and WebGestalt were adjusted as FDR using the Benjamini-Hochberg Method, with the statistical significance threshold set as FDR<0.05. GO functional annotation terms was presented by their Enrichment scores, calculated by -log10^[Pvalue]^, and over-representation analysis on WebGestalt was displayed by presenting each signalling pathways’ enrichment ratios and FDR.

## RESULTS

### Acidified PBS used as a vehicle alters the transcriptome of endometrial epithelial cells exposed in a microfluidic device

Differentially expressed genes (DEGs) from each of the three treatment groups (control, vehicle control [VC], and insulin) were compared with one another to minimise the confounder of VC added in with insulin. Comparison of control to VC altered the expression of 22 transcripts in total (Figure 2: Table 1). Genes involved in the biological processes of Mitochondrial electron transport, NADH to ubiquinone and Mitochondrial respiratory chain complex I assembly were overrepresented in the list of DEGs (Table 2). There were more transcripts associated with the cellular components of Respiratory chain, Mitochondrial inner membrane, Mitochondrial respiratory chain complex I, and Mitochondrion as well as the molecular functions of NADH dehydrogenase (ubiquinone) activity and Cytochrome-c oxidase activity, than one would have expected by chance (For full details see Table 2). While there were a number of overrepresented pathways (including those associated with thiamine metabolism, and folate biosynthesis) none had an FDR of <0.05 (Figure 3: Table 3).

**Table 1.** Transcript ID and description, associated with differentially expressed transcripts between control, vehicle control, and Insulin treated groups. DEGs were identified following RNA sequencing of cells treated with control, vehicle control (acetic acid), or Insulin (2mg/mL) exposed endometrial epithelial (Ishikawa) cells cultured in a microfluidics device for 24 hr (n=3 biological replicates).

| Control v Vehicle Control |  | Vehicle Control v Insulin |  |
| --- | --- | --- | --- |
| Transcript | Description | Transcript | Description |
| **MT-CYB** | Mitochondrially encoded cytochrome B | **SSB** | Small RNA binding exonuclease protection factor La |
| **MT-CO1** | Mitochondrially encoded cytochrome C oxidase 1 | **SPCS1** | Signal peptidase complex subunit 1 |
| **MT-CO2** | Mitochondrially encoded cytochrome C oxidase 2 | **BTF3** | Basic transcription factor 3 |
| **MT-ND3** | Nicotinamide adenine dinucleotide + hydrogen (NADH)-Ubiquinone oxidoreductase Chain 3 | **GGCT** | Gamma-glutamyl cyclotransferase |
| **MT-ND5** | NADH-Ubiquinone oxidoreductase chain 5 | **RPL7A** | Ribosomal protein L7a |
| **COA6** | Cytochrome C oxidase assembly factor 6 | **RAPSN** | Receptor associated protein of the synapse |
| **ALPG** | Alkaline phosphatase, germ cell | **HTATSF1** | HIV-1 Tat specific factor 1 |
| **THEGL** | Theg spermatid protein like | **TMSB10** | Thymosin beta 10 |
| **HIST1H2BN** | Histone H2B type 1-N | **RPS6** | Ribosomal protein S6 |
| **HMGA1** | High mobility group AT-Hook 1 |  |  |
| **SCX** | Scleraxis BHLH transcription factor |  |  |
| **MKI67** | Marker of proliferation Ki-67 |  |  |
| **RPL36AL** | Ribosomal protein L36A-like |  |  |
| **RPL19** | Ribosomal protein L19 |  |  |
| **PRKACA** | Protein kinase cAMP-activated catalytic subunit alpha |  |  |
| **CTSZ** | Cathepsin Z |  |  |
| **PPDPF** | Pancreatic progenitor cell differentiation and proliferation factor |  |  |
| ***TCEAL4*** | Transcription elongation factor A like 4 |  |  |
| ***MT-ND1*** | Mitochondrially encoded NADH: ubiquinone oxidoreductase core subunit 1 |  |  |
| ***MT-ND4*** | Mitochondrially encoded NADH: ubiquinone oxidoreductase core subunit 4 |  |  |
| ***TMSB10*** | Thymosin beta 10 |  |  |
| ***RPS6*** | Ribosomal protein S6 |  |  |

**Table 2.**
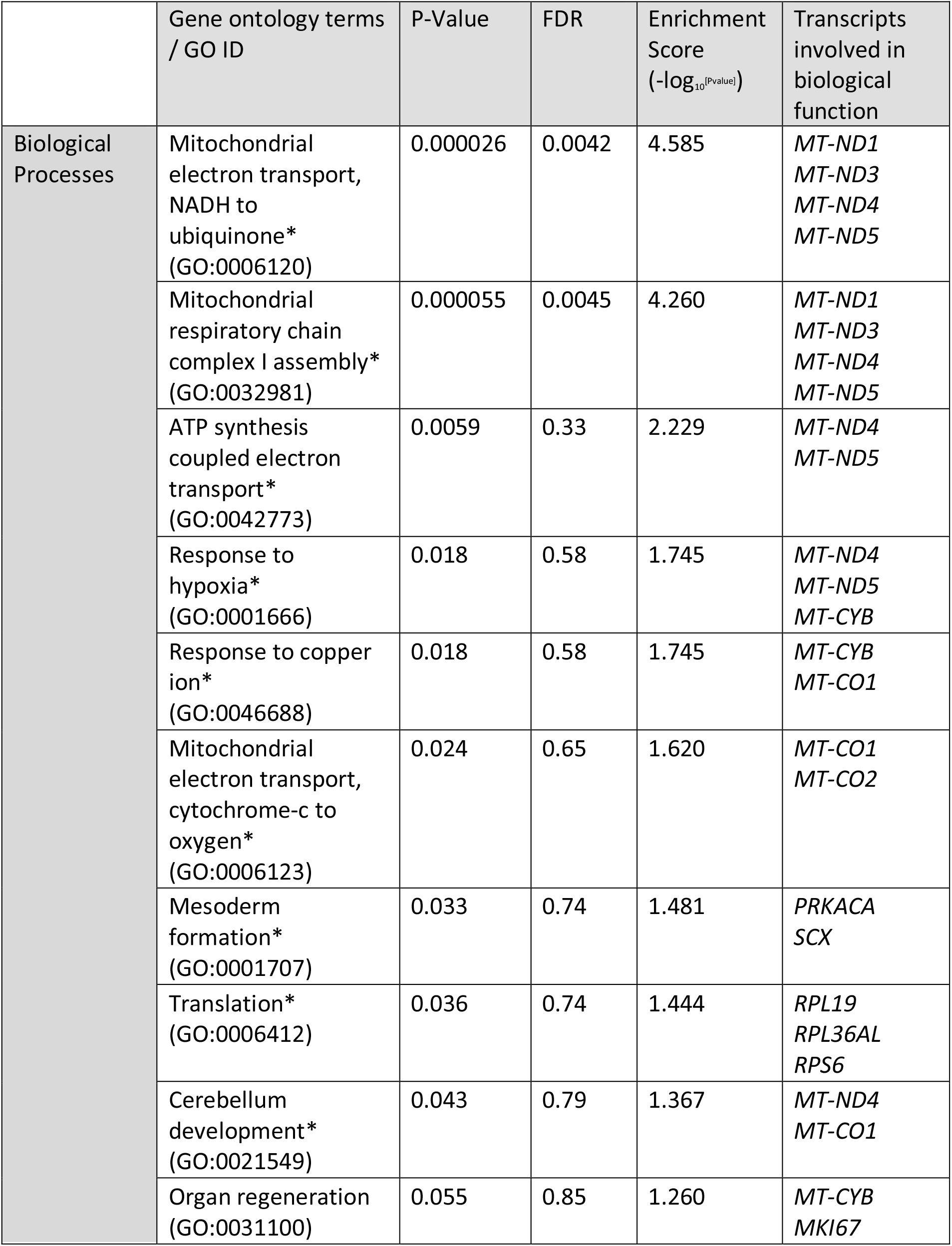
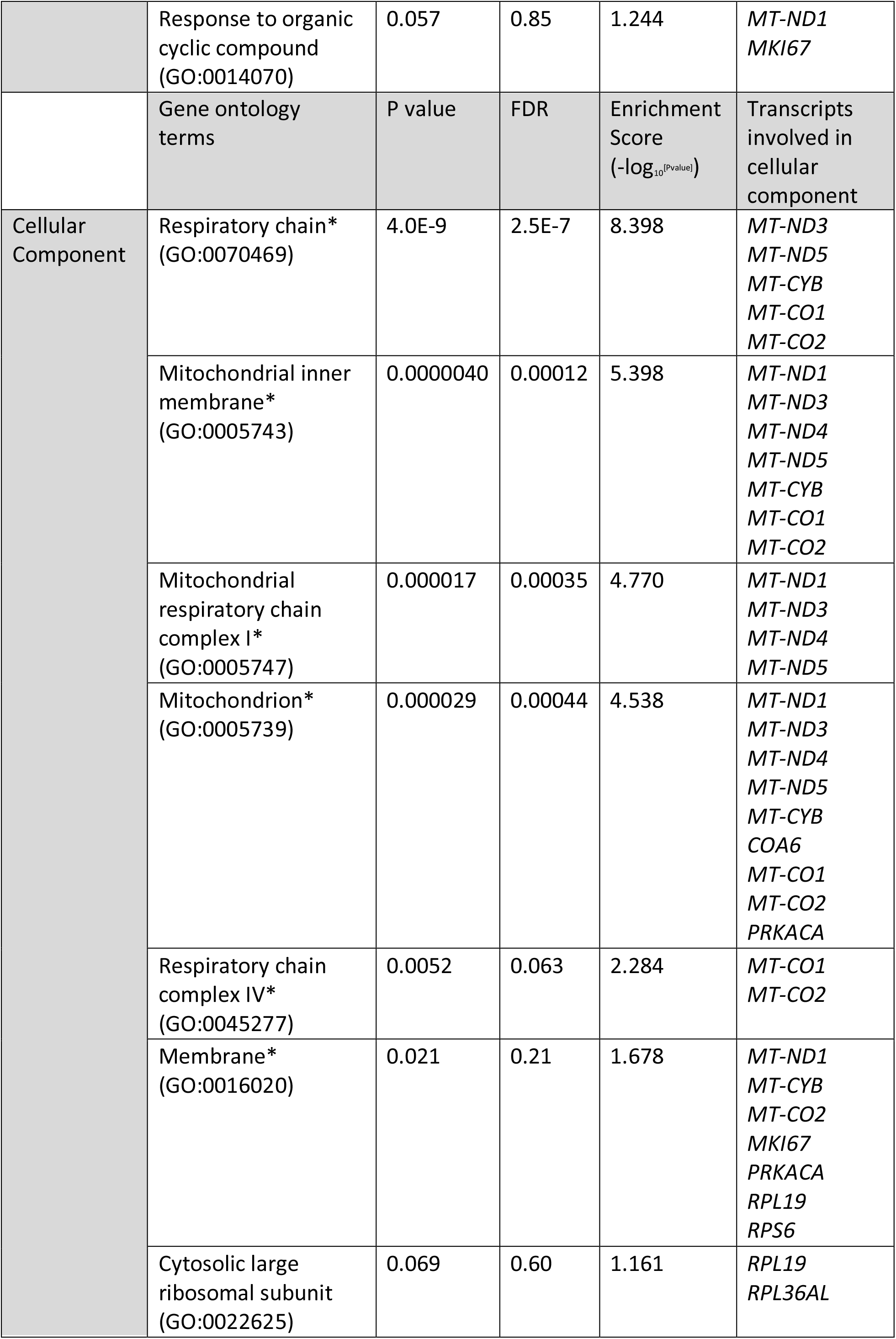
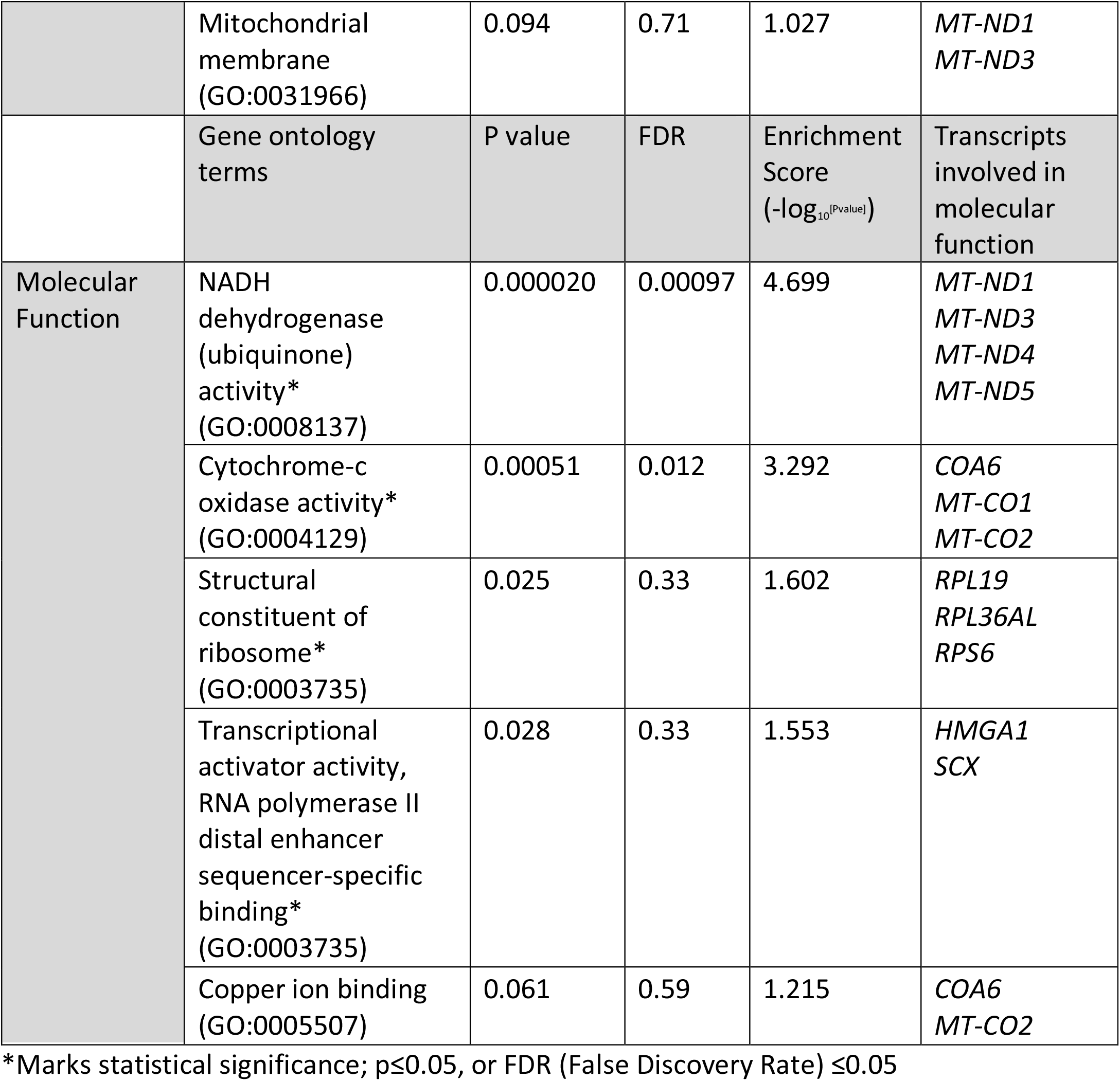
Gene Ontology terms involved in biological processes, cellular component, and molecular function associated with differentially expressed transcripts between control v vehicle control groups. Overrepresented GO terms and their descriptions associated with DEGs identified between control and vehicle exposed Ishikawa cells cultured in a microfluidics device for 24 hr (n=3 biological replicates). DEGs were identified following RNA sequencing of cells and overrepresentation analysis determined using Webgestelt.

**Table 3.**
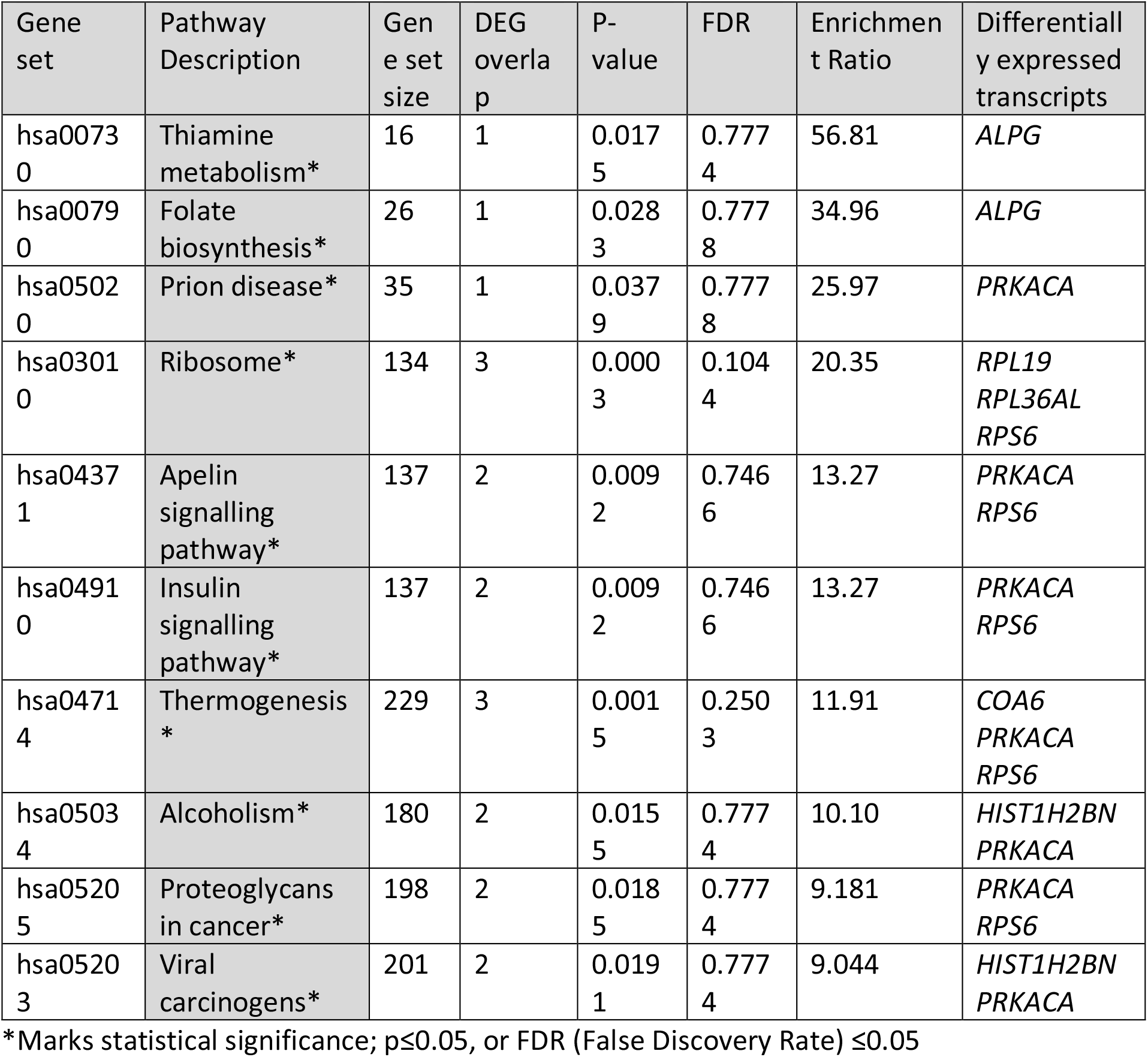
Overrepresented signalling pathways in DEGs between control v vehicle control groups. Overrepresented signalling pathways and their descriptions associated with DEGs identified between control and vehicle exposed Ishikawa cells cultured in a microfluidics device for 24 hr (n=3 biological replicates). DEGs were identified following RNA sequencing of cells and overrepresentation analysis determined using Webgestelt.

**Figure 2.**
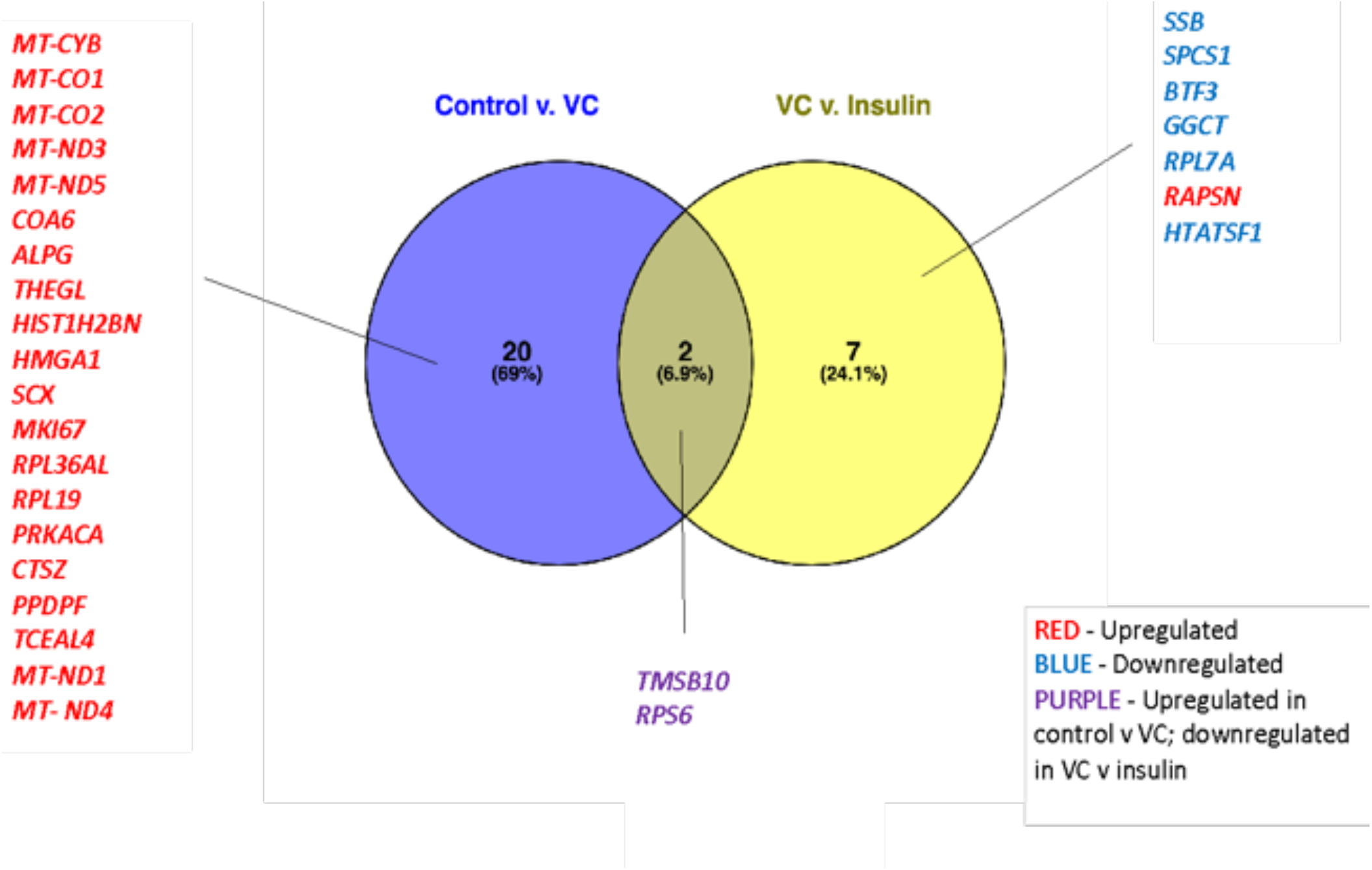
Venn diagram of DEGs identified in endometrial epithelial cells following exposure to control media, vehicle, or Insulin in a microfluidic device for 24 hrs. Ishikawa cells (n=3) were expose to control medium, vehicle control (acetic acid: VC), or Insulin (2 mg/mL) for 24 hrs in a microfluidics device (1 ul/min). Following RNA sequencing DEGs in treatment groups were identified using a cut off of p<0.05 and log2FoldChange >0 or log2FoldChange <0; fold change differences for transcripts were determined using paired t-tests. “Control v VC” demonstrates the transcripts altered when exposing Ishikawa cells to Control compared to those altered in VC. “VC v insulin” shows the comparison of DEGs expressed when Ishikawa cells were exposed to VC and insulin treatment. Results were found to be significant when p<0.05, and the results were derived from n=3 biological replicates.

**Figure 3.**
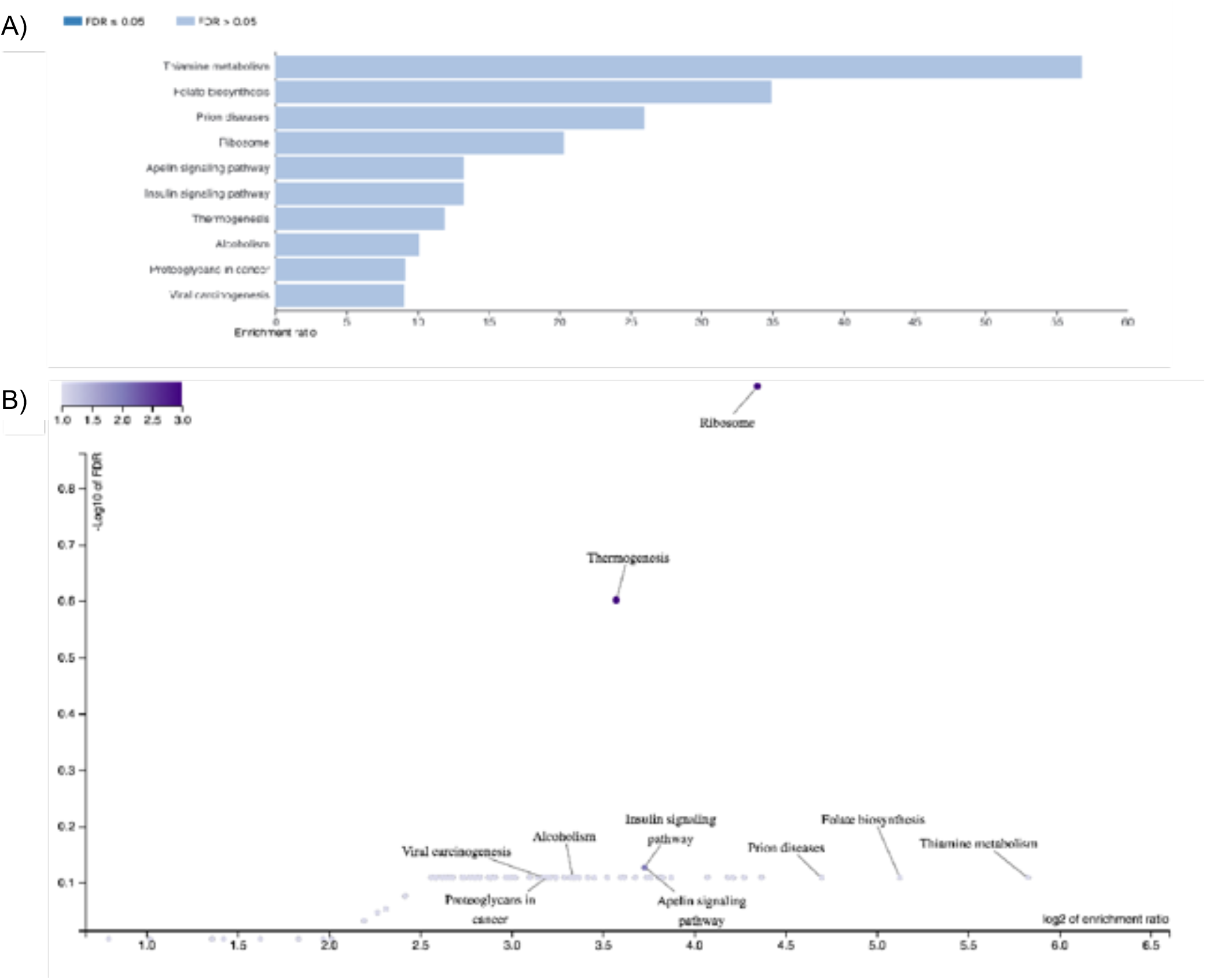
Over-representation analysis (ORA) graphs depicting enriched KEGG pathways associated DEGs found in Control v Vehicle Control groups. ORA was produced by Webgestalt with the gene identifier “Gene Symbol” and functional database of “KEGG pathways”. Benjamini-Hochberg method was used to adjust P-values and presented as FDR, FDR≤0.05 being the threshold for statistical significance. A) Bar chart illustrates enrichment ratio plotted against enriched pathway identified through ORA, with the blue-colour shade indicating statistical significance based on FDR (dark blue = FDR ≤0.05; light blue = FDR>0.05). The chart is organised according to decreasing enrichment ratio, with the top pathway being most enriched and bottom being least enriched. B) Volcano plot plotting -log10 of FDR against log2 of enrichment ratio for each pathway. Darker purple shade of plotted dots indicates pathway of interest, as it demonstrates notable magnitude in both log2 of enrichment ratio and -log10 of FDR compared to other regulatory networks.

### Exposure of endometrial epithelial cells to Insulin in a microfluidics device altered the transcriptional profile

Exposure of Ishikawa cells to Insulin altered the expression of 9 genes (Figure 2: Table 1). Two of these transcripts were altered in both comparisons however, *TSMB10* and *RPS6* were increased in expression in control versus VC but were significantly decreased in VC compared to insulin treated cells. The molecular function of Poly(A) RNA binding was significantly overrepresented in cells exposed to insulin in a microfluidic device (Figure 4: Table 4). While genes associated with the pathways Protein export, Glutathione metabolism, and Ribosome (Table 5) were significantly overrepresented following insulin exposure. Comparisons of the overrepresented biological processes, cellular components, and molecular functions are provided in Figure 5.

**Table 4.** Gene Ontology terms involved in biological processes, cellular component, and molecular function associated with differentially expressed transcripts between vehicle control v Insulin treated groups. Overrepresented GO terms and their descriptions associated with DEGs identified between vehicle and Insulin (2 mg/ml) exposed Ishikawa cells cultured in a microfluidics device for 24 hr (n=3 biological replicates). DEGs were identified following RNA sequencing of cells and overrepresentation analysis determined using Webgestelt.

| | Gene ontology terms | P-Value | FDR | Enrichment Score<br>($-\log_{10}^{[Pvalue]}$) | Transcripts involved in biological function |
| --- | --- | --- | --- | --- | --- |
| Biological Function | SRP-dependent cotranslational protein targeting to membrane* (GO:0006614) | 0.044 | 0.82 | 1.357 | *RPL7A*<br>*RPS6* |
|  | Viral transcription (GO:0019083) | 0.052 | 0.82 | 1.284 |  |
|  | Nuclear-transcribed mRNA catabolic process, nonsense-mediated decay (GO:0000184) | 0.055 | 0.82 | 1.260 |  |
|  | Translational initiation (GO:0006413) | 0.063 | 0.82 | 1.201 |  |
|  | rRNA processing (GO:0006364) | 0.098 | 0.99 | 1.009 |  |
| | Gene ontology terms | P-value | FDR | Enrichment Score<br>($-\log_{10}^{[Pvalue]}$) | Transcripts involved in cellular component |
| Cellular Component | Intracellular ribonucleoprotein complex (GO:1990904) | 0.58 | 1.0 | 0.2366 | *SSB*<br>*RPS6* |
|  | Ribosome (GO:0005840) | 0.71 | 1.0 | 0.1487 | *RPL7A*<br>*RPS6* |
| | Gene ontology terms | P-value | FDR | Enrichment Score<br>($-\log_{10}^{[Pvalue]}$) | Transcripts involved in molecular function |
| Molecular Function | Poly(A) binding* (GO:0008143) | 0.0011 | 0.021 | 2.959 | *HTATSF1*<br>*SSB*<br>*BTF3*<br>*RPL7A*<br>*RPS6* |
|  | RNA binding* (GO:0003723) | 0.026 | 0.24 | 1.585 | *HTATSF1*<br>*SSB* |
|  |  |  |  |  | *RPL7A* |

\*Marks statistical significance; $p \leq 0.05$ , or FDR (False Discovery Rate) $\leq 0.05$

**Table 5.** Overrepresented signalling pathways in DEGs between control v vehicle control groups. Overrepresented signalling pathways and their descriptions associated with DEGs identified between vehicle and Insulin (2mg/mL) exposed Ishikawa cells cultured in a microfluidics device for 24 hr (n=3 biological replicates). DEGs were identified following RNA sequencing of cells and overrepresentation analysis determined using Webgestelt.

| Gene set | Pathway Description | Gene set size | DEG overlap | FDR | P-value | Enrichment Ratio | Differentially expressed transcripts involved |
| --- | --- | --- | --- | --- | --- | --- | --- |
| Hsa03060 | Protein export* | 23 | 1 | 1 | 0.0157 | 63.23 | *SPCS1* |
| Hsa00480 | Glutathione metabolism* | 56 | 1 | 1 | 0.0379 | 25.97 | *GGCT* |
| Hsa03010 | Ribosome* | 134 | 2 | 1 | 0.0033 | 21.70 | *RPL7A*<br>*RPS6* |
| Hsa01521 | EGFR tyrosine kinase inhibitor resistance | 79 | 1 | 1 | 0.0532 | 18.41 | *RPS6* |
| Hsa04066 | HIF-1 signalling pathway | 100 | 1 | 1 | 0.0669 | 14.54 | *RPS6* |
| Hsa05322 | Systemic Lupus Erythematosus | 133 | 1 | 1 | 0.0882 | 10.93 | *SSB* |
| Hsa04371 | Apelin signalling pathway | 137 | 1 | 1 | 0.0908 | 10.62 | *RPS6* |
| Hsa04910 | Insulin signalling pathway | 137 | 1 | 1 | 0.0908 | 10.62 | *RPS6* |
| Hsa04150 | mTOR signalling pathway | 151 | 1 | 1 | 0.0996 | 9.631 | *RPS6* |
| Hsa05205 | Proteoglycans in cancer | 198 | 1 | 1 | 0.1290 | 7.344 | *RPS6* |

\*Marks statistical significance; $p \leq 0.05$ , or FDR (False Discovery Rate) $\leq 0.05$

**Figure 4.**
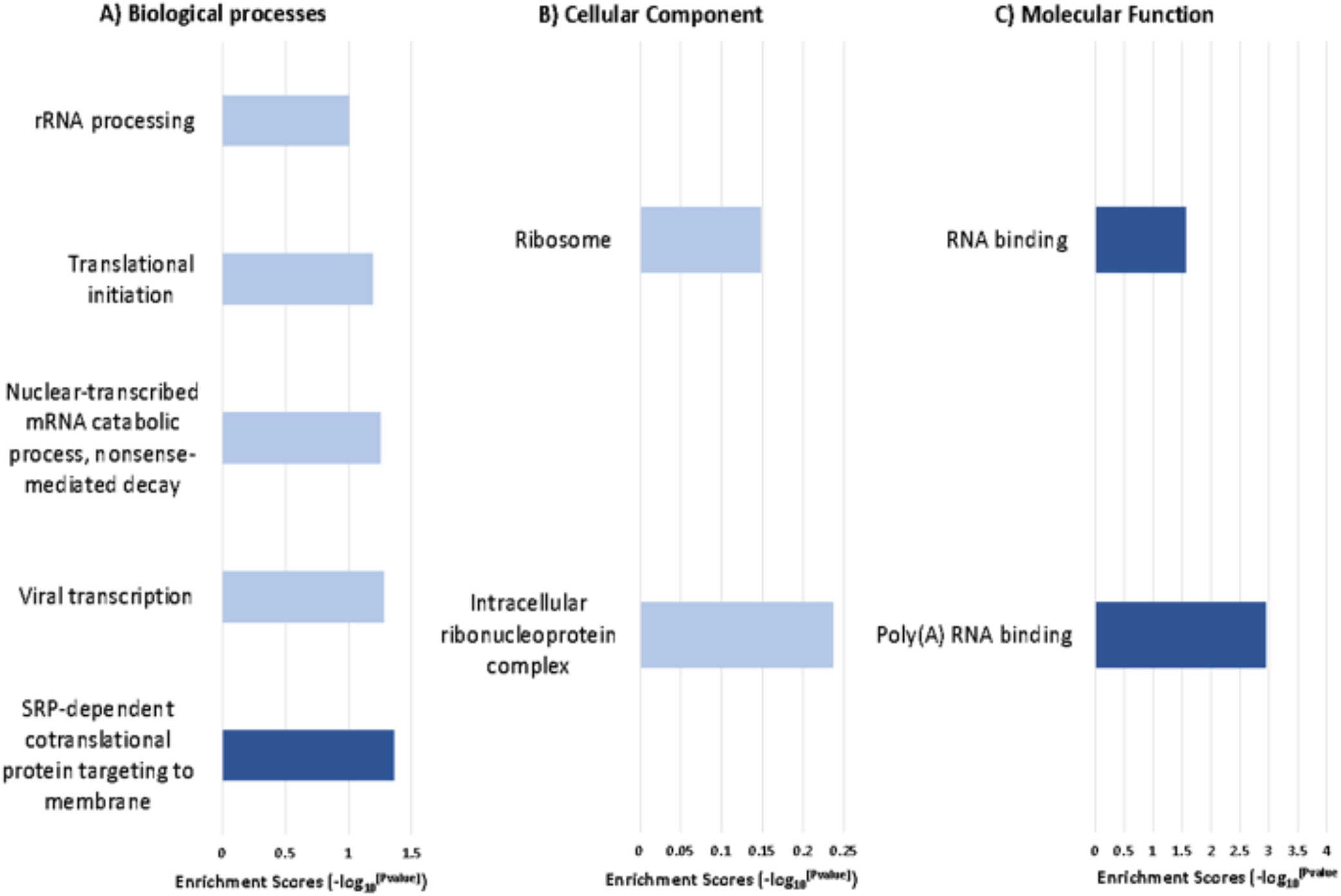
Over-representation analysis (ORA) graphs depicting enriched KEGG pathways associated DEGs found in Vehicle Control v Insulin treated cells. ORA were produced by Webgestalt with the gene identifier “Gene Symbol” and functional database of “KEGG pathways”. Benjamini-Hochberg method was used to adjust P-values and presented as FDR, FDR≤0.05 being the threshold for statistical significance. A) Bar chart illustrates enrichment ratio plotted against enriched pathway identified through ORA, with the blue-colour shade indicating statistical significance based on FDR (dark blue = FDR ≤0.05; light blue = FDR>0.05). The chart is organised according to decreasing enrichment ratio, with the top pathway being most enriched and bottom being least enriched. B) Volcano plot plotting -log10 of FDR against log2 of enrichment ratio for each pathway. Darker purple shade of plotted dots indicates pathway of interest, as it demonstrates notable magnitude in both log2 of enrichment ratio and -log10 of FDR compared to other regulatory networks.

**Figure 5.**
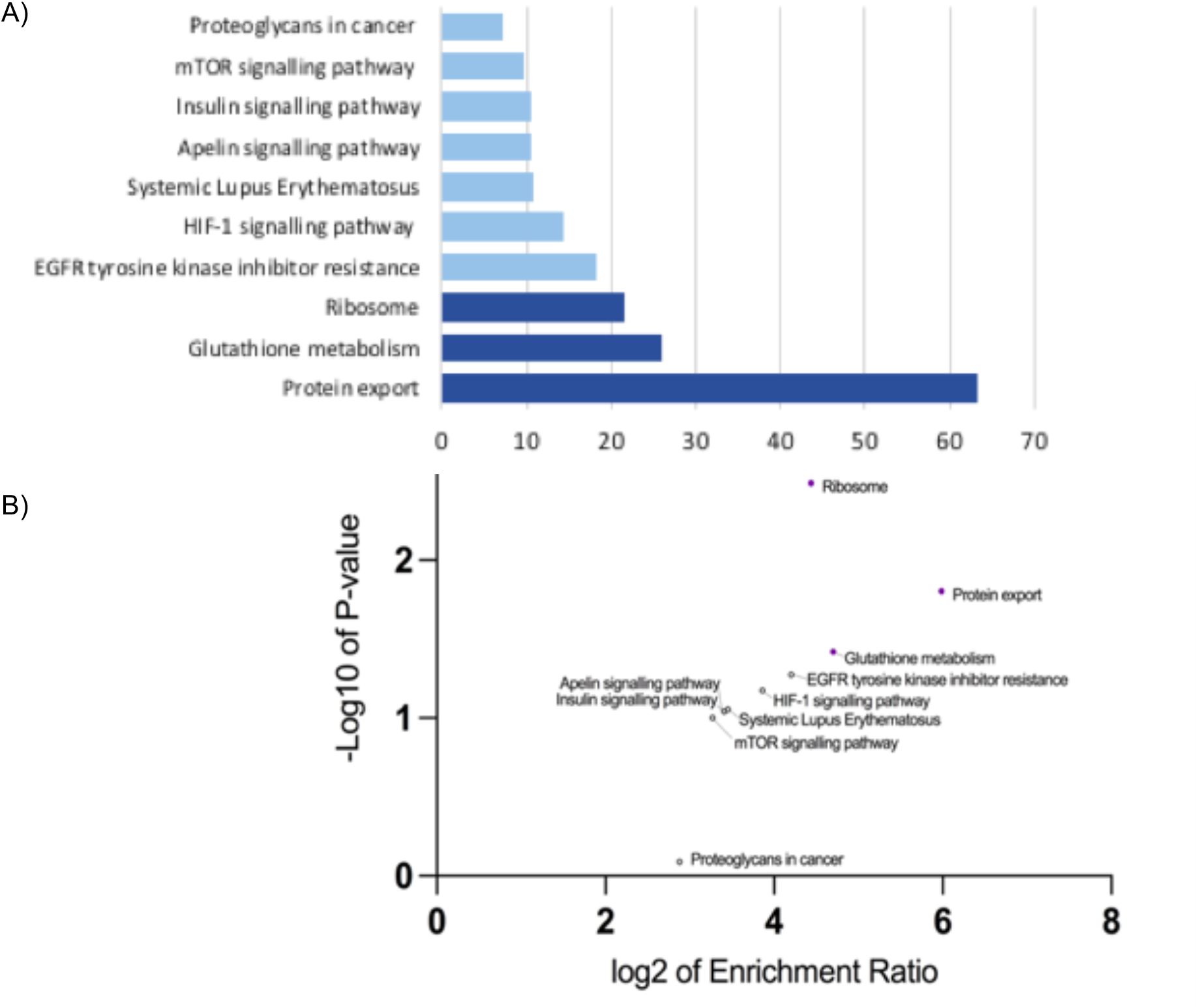
Over-representation analysis (ORA) graphs depicting enriched KEGG pathways associated DEGs found in Vehicle Control v Insulin groups. ORA were produced by Webgestalt with the gene identifier “Gene Symbol” and functional database of “KEGG pathways”. Benjamini-Hochberg method was used to adjust P-values and presented as FDR, FDR≤0.05 being the threshold for statistical significance. A) Bar chart illustrates enrichment ratio plotted against enriched pathway identified through ORA, with the blue-colour shade indicating statistical significance based on FDR (dark blue = FDR ≤0.05; light blue = FDR>0.05). The chart is organised according to decreasing enrichment ratio, with the top pathway being most enriched and bottom being least enriched. B) Volcano plot plotting -log10 of FDR against log2 of enrichment ratio for each pathway. Darker purple shade of plotted dots indicates pathway of interest, as it demonstrates notable magnitude in both log2 of enrichment ratio and -log10 of FDR compared to other regulatory networks.

**Figure 6.**
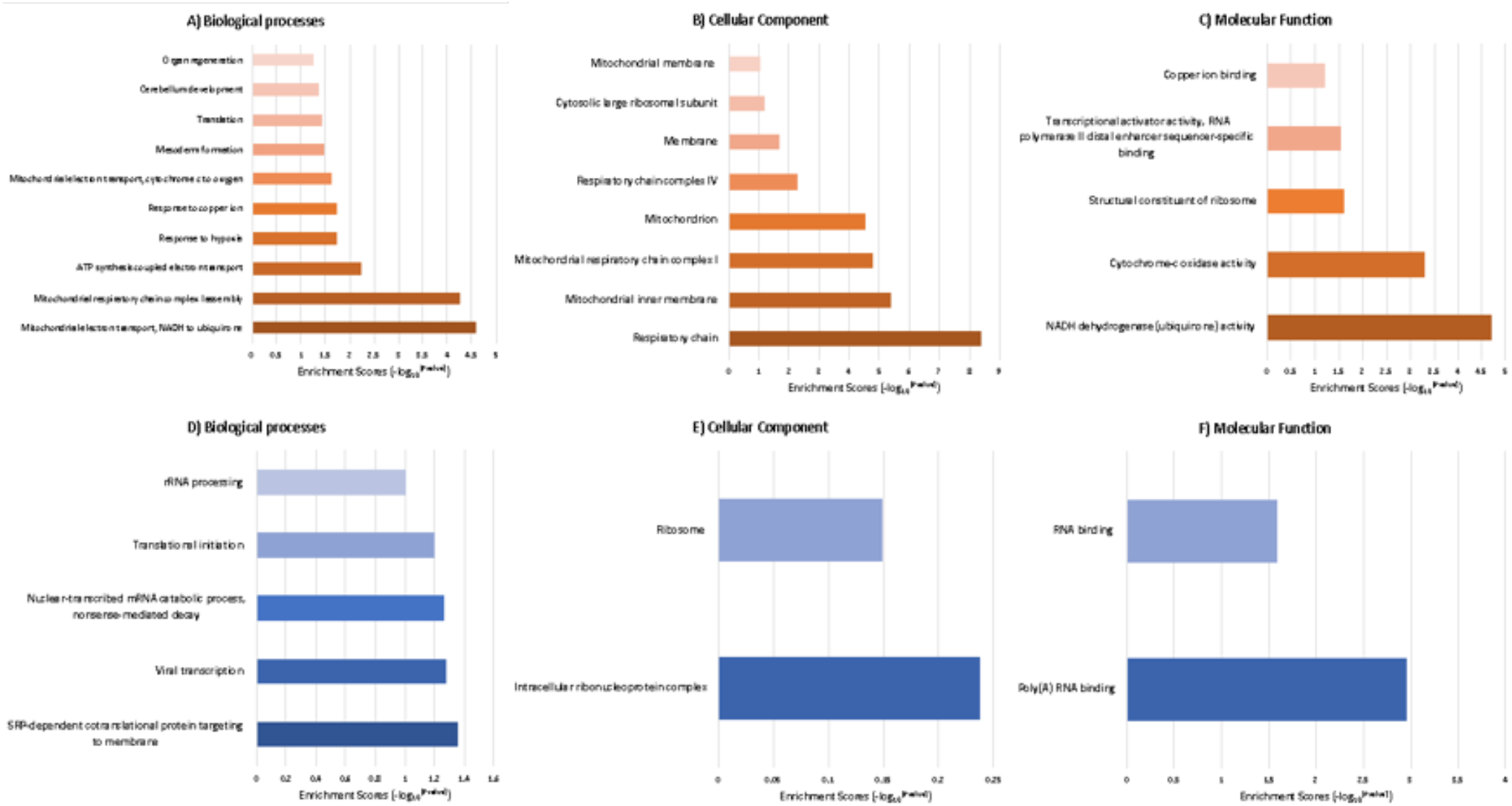
Bar charts indicating overrepresented GO terms for biological processes, cellular component, and molecular functions associated with DEGs. Charts are divided into two groups: A) biological processes, B) cellular component, and C) molecular function for n=22 DEGs involved in Control v. VC group, and D, E, and F depicting gene ontology terms found in from n=9 DEGs involved in VC v. Insulin group. All DEGs involved in GO terms in A), B) and C) were upregulated, thus shown in orange. All DEGs involved in the GO terms in D), E), and F) were downregulated as RAPSN (the only upregulated DEG out of n=9 DEG in VC v Insulin) was not associated with any of the functional annotations, thus shown in blue. GO terms and p-values were determined by overrepresented functional annotation analysis using DAVID. P-values were used to calculate enrichment scores (-log10[Pvalue]). Colour gradient depicts size of enrichment score, darker being more enriched (or having higher p-value), and lighter being less enriched.

## DISCUSSION

To the best of our knowledge, our study is the first to use a microfluidics approach to mimic exposure of the endometrial epithelium to obsesogenic concentrations of insulin associated with maternal circulation to enhance our understanding of the mechanism by which endometrial function may be affected obesity. This was conducted by allowing exposure of Ishikawa cells in a microfluidics device to insulin concentration associated with an obese environment and identifying the associated pathways and functional annotations that were modified. Our results indicate that maternal metabolic stressors, insulin, may alter uterine function and receptivity by altering specific transcripts in the endometrial epithelium and propose the mechanisms by which this may occur (Figure 7).

**Figure 7.**
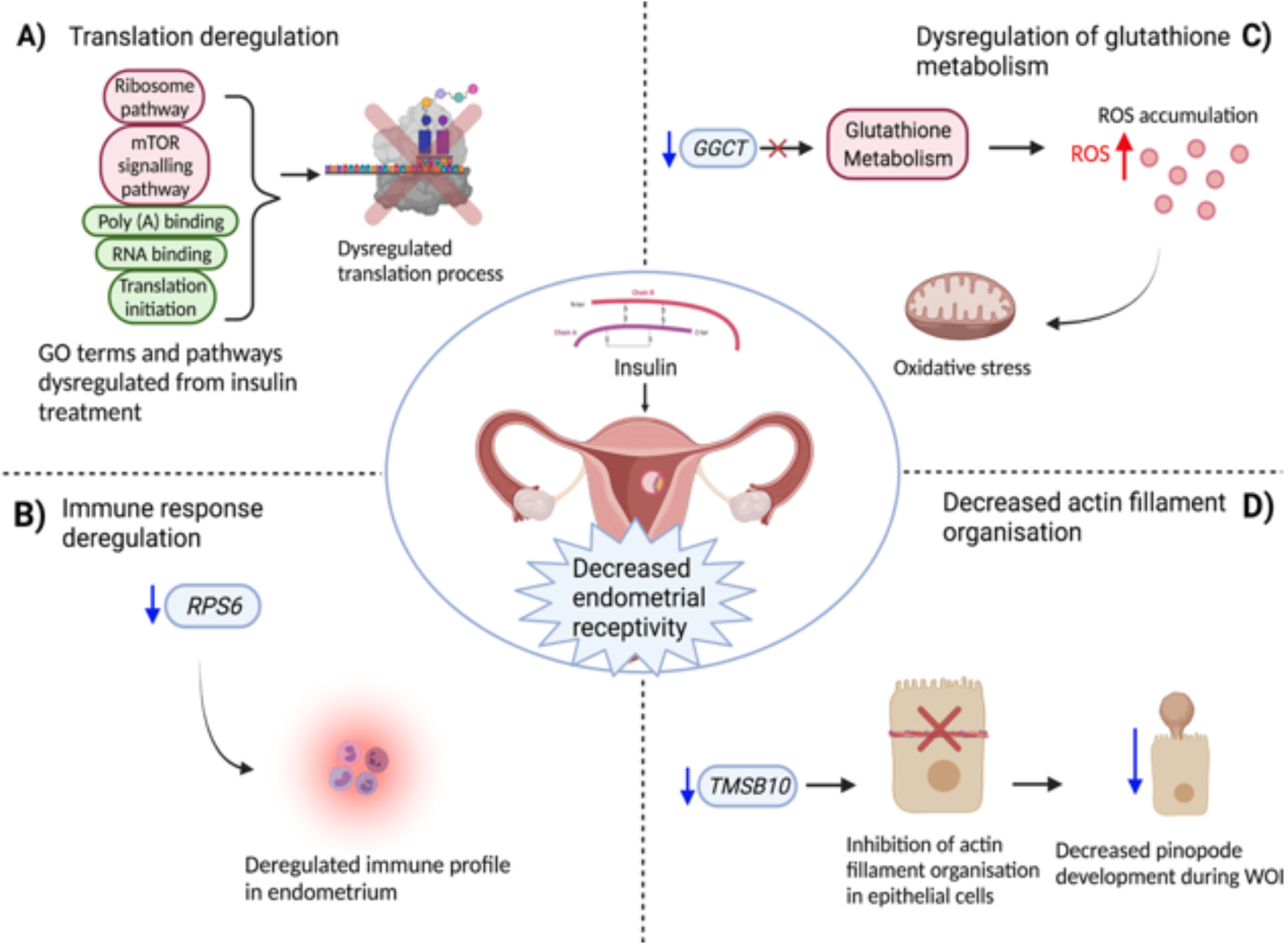
Proposed mechanism by which insulin alters endometrial function. Created using Biorender.

The vehicle control used to deliver Insulin to the cells altered biological functions of mitochondrial electron transport (NADH to ubiquinone), mitochondrial respiratory chain complex I assembly, ATP synthesis; molecular functions of NADH dehydrogenase (ubiquinone activity), and cytochrome-c oxidase activity were altered. While NADH dehydrogenase promotes the smooth functioning of respiration and ATP synthesis by stimulating the assembly of respiratory chain complex I, they may have an additional role of producing apoptosis-inducing fragments to promote cell death [28, 29]. This alternative role of NADH dehydrogenase in our findings with VC can be further substantiated by the fact acetic acid is reported to induce apoptosis in a gastric cell line [30]. Apoptosis in cells is initiated by acetic acid by mitochondrial dysfunction, cytochrome-c release, and along with reactive oxygen species accumulation, which seem to correlate with the GO terms found in control v VC [31]. Using acetic acid might have stimulated mitochondria-induced apoptosis in the Ishikawa cells.

The processes and signalling pathways modified by insulin treatment may provide a major insight into the function of insulin endometrial receptivity specifically those related to processes involving translation. Translation initiation, is facilitated by binding of ribosomes to the 5’cap of mRNAs and is mediated by the eukaryotic translation initiation factor 4F (eIF4F) complex. One of the major components of the eIF4F complex is the eIF4G, which aids in recruiting and binding of the ribosome. The 3’ poly(A) tail end of the mRNA binds to the poly(A) binding protein (PABP) with PABP and elF4G interaction synergistically initiates translation [32, 33]. The mTOR signalling pathway was also shown to be modified by insulin in our study and this signalling pathway is implicated in translational initiation by phosphorylating translation factors such as EIF4E binding protein [34]. Furthermore, *RPS6* phosphorylation (a DEG significantly downregulated by insulin in our findings) within the mTOR pathway is widely accepted to stimulate protein synthesis and translation [35]; cellular protein synthesis capacity is widened by the mTOR pathway and is critical to cell survival [36].

Translation modulation, and increased ribosome synthesis are implicated in endometrial epithelial cell function and receptivity given the dynamic nature of the human endometrium [32, 37]. A recent study identified the proteins involved in repeated implantation failure revealed were involved in translation pathways, posttranslational modification and ribosomal pathways were heavily enriched in the differentially expressed proteins [38]. It is thus clear that our findings show evidence that insulin treatment potentially modifies the translational profile of endometrial epithelial cells.

Insulin treatment induced differential expression of transcripts from Ishikawa cells, which may provide insight into how obesity-related insulin changes can alter endometrial function. *RPS6 (*Ribosomal protein S6) is a transcript that is associated with many of the enriched and over-represented pathways in analysis. The gene is highly expressed in the endometrium, with 1752 Transcripts per million expressed in the endometrial tissue [39]. Aside from its role in translation regulation in the ribosomal and mTOR pathway, it is also implicated in immune response regulation [39]. Phosphorylation of *RPS6* is necessary for T-cell differentiation in the thymus [40], plays a key role in the mTOR pathway, which has been demonstrated to be involved in T-cell signalling [41]. The immune profile of the endometrium plays a critical role in implantation success via T-cell mediated immune tolerasiation [42]. In fact, research showed that patients experiencing RIF during IVF had a dysregulated endometrial uterine immune profile [43]. Taking into consideration of the well-documented responsibility of *RPS6* in regulating T cell expression during the immune response, and that insulin downregulated *RPS6* expression, it can be hypothesised that high circulating insulin in the endometrium may compromise the uterine immune profile and consequent receptivity.

*GGCT* encodes an enzymatic protein essential for glutathione metabolism. Dysregulation in the glutathione metabolism pathway is associated with insulin resistance, hyperinsulinaemia, and type 2 diabetes [44, 45]. It is thought that GGCT is important in cellular protective mechanisms by salvaging glutathione, thereby providing an antioxidant effect to cells [46]. Glutathione is a critical modulator of normal cellular function such as gene expression, protein synthesis, and immune response but is primarily involved in antioxidative roles and nutrient metabolism [47]. Glutathione deficiency leads to accumulation of free-radicals and oxidative stress [48]. Glutathione metabolism also may play a major role in fertility and endometrial function, as Xu et al. [49] demonstrated that this pathway is crucial in reducing hydrogen peroxide (a type of reactive oxygen species that causes oxidative stress) during decidualisation. High concentrations of reactive oxygen species (ROS) and oxidative stress adversely affects implantation process, early embryo development, and can eventually cause implantation failure [50] as well as normal endometrial function, making the environment unsuitable to support growth and development of the embryo [51]. We propose that insulin can disturb glutathione homeostasis in the endometrium, thereby leading to oxidative stress and subsequent effects on uterine receptivity to implantation.

The downregulation of *TMSB10* with insulin treatment encodes a protein that aids the processes of actin filament organization and actin binding [39]. During the window of receptivity, the human endometrial surface goes through ultrastructural modifications which includes the development of pinopodes, which is tightly managed by cytoskeletal actin filaments [52, 53]. Although previously controversial, recent literature has deemed the presence of pinopodes in the endometrial epithelia as a credible biomarker of uterine receptivity [52, 54]. Filamentous actin expression coincides with pinopode formation during receptivity on the apical surface of luminal epithelial cells [55]. In the endometria of from those suffering recurrent pregnancy loss, both ezrin and thrombomodulins were downregulated which disrupted the organisation of cytoskeletal actin filaments and may ultimately impaired pinopode formation. *TMSB10* was differentially expressed and highly responsive to oestradiol and progesterone [56] and has been associated with a functional and receptive endometrium [57]. We conclude that insulin concentrations can downregulate *TMSB10*, which may contribute to impaired implantation capacity of the endometrium.

Despite numerous reports on the potential contribution of insulin as a metabolic stressor in obesity, the mechanism by which uterine receptivity may be compromised is still unknown. Existing literature studying this phenomenon and underlying mechanisms utilised hyperinsulinaemic animal models [17, 58], which may not accurately reflect the inner workings of a human endometria. Our findings aimed to provide insight into this gap in knowledge through using a microfluidics approach. Microfluidics can also simulate shear stress and communications between cells, elucidating that the constant flow of maternal circulation within the endometria can be modelled effectively through the device [59]. In conclusion, our results indicate that insulin alters the transcriptional profile of the endometrial epithelial cells when cultured in a microfluidics device. The biological processes and signalling pathways associated with differentially expressed transcripts found following insulin treatment were related to certain mechanisms of endometrial receptivity and implantation (Figurer 7). Specifically, genes and processes related translation, immune response regulation, glutathione metabolism, and actin filament organisations seem to be dysregulated with insulin treatment, which may ultimately impair implantation success and reduce pregnancy success in those suffering from obesity.

## ACKNOWLEDGEMENTS

NF’s lab is supported by funding from N8 agri-food pump priming, QR GCRF, UN-CRP, as well as BBSRC grant number BB/R017522/1. RNA sequencing analysis was undertaken on ARC3, part of the High-Performance Computing facilities at the University of Leeds, UK. Figures 1 and 7 were created using Biorender.

## Notes

### Competing Interest Statement

The authors have declared no competing interest.

